# The relaxin receptor Lgr3 mediates growth coordination and developmental delay during *Drosophila melanogaster* imaginal disc regeneration

**DOI:** 10.1101/032508

**Authors:** Jacob S. Jaszczak, Jacob B. Wolpe, Rajan Bhandari, Rebecca G. Jaszczak, Adrian Halme

## Abstract

Damage to *Drosophila melanogaster* imaginal discs activates a regeneration checkpoint that 1) extends larval development and 2) coordinates the regeneration of the damaged disc with the growth of undamaged discs. These two systemic responses to damage are both mediated by Dilp8, a member of the insulin/IGF/relaxin family of peptide hormones, which is released by regenerating imaginal discs. Growth coordination between regenerating and undamaged imaginal discs is dependent on Dilp8 activation of NOS in the prothoracic gland (PG), which slows the growth of undamaged discs by limiting ecdysone synthesis. Here we demonstrate that the *Drosophila* relaxin receptor homologue Lgr3, a leucine-rich repeat-containing G-protein coupled receptor, is required for Dilp8-dependent growth coordination and developmental delay during the regeneration checkpoint. Lgr3 regulates these responses to damage via distinct mechanisms in different tissues. Using tissue-specific RNAi disruption of *Lgr3* expression, we show that Lgr3 functions in the PG upstream of nitric oxide synthase (NOS), and is necessary for NOS activation and growth coordination during the regeneration checkpoint. When Lgr3 is depleted from neurons, imaginal disc damage no longer produces either developmental delay or growth inhibition. To reconcile these discrete tissue requirements for Lgr3 during regenerative growth coordination, we demonstrate that Lgr3 activity in the both the CNS and PG is necessary for NOS activation in the PG following damage. Together, these results identify new roles for a relaxin receptor in mediating damage signaling to regulate growth and developmental timing.

## Introduction

Growth rate and developmental time must be regulated in concert to ensure that organs develop to the correct size and proportion. Following damage to imaginal discs, *Drosophila* larvae activate a regeneration checkpoint that delays development and slows the growth of undamaged imaginal discs. These systemic responses to damage may function to coordinate regeneration with the growth and development of undamaged tissues (Stieper *et al*. 2008; Halme *et al*. 2010; Parker and Shingleton 2011; Jaszczak *et al*. 2015). The peptide Dilp8 is required for both delay and growth coordination and is secreted by regenerating imaginal discs to activate the regeneration checkpoint (Colombani *et al*. 2012; Garelli *et al*. 2012). Dilp8 induces developmental delay by inhibiting production of the neuropeptide prothoracicotropic hormone (PTTH) in the central nervous system (CNS) (Halme *et al*. 2010; Colombani *et al*. 2012), whereas Dilp8 inhibits growth of the undamaged imaginal discs by reducing biosynthesis of the steroid hormone ecdysone through activation of nitric oxide synthase (NOS) in the prothoracic gland (PG)(Jaszczak *et al*. 2015).

Dilp8 has been classified as a member of the insulin/IGF/relaxin family of peptide hormones (Garelli *et al*. 2012). Relaxin receptors in mammals belong to a larger family of leucine-rich repeat-containing G-protein coupled receptors (LGRs), which are subdivided into type A vertebrate gonadotropin receptors, type B Wnt agonist R-spondin receptors Lgr4/5/6, which also includes the *Drosophila* bursicon receptor (*Lgr2/rickets*), and type C relaxin receptors (Barker *et al*. 2013). The different classes of LGR receptors are distinguished by different numbers of extracellular leucine-rich repeats (LRRs), the presence of a low-density lipoprotein receptor class A domain (LDLa), and the structure of the hinge region connecting the transmembrane region to the LRR domain. Here we demonstrate that the relaxin receptor Lgr3 mediates Dilp8 signaling during the regeneration checkpoint developmental delay and growth coordination. We find that Lgr3 functions in the CNS as well as in the PG to regulate the coordination of growth and that these two Lgr3 pathways converge on the regulation of NOS activation in the PG.

## Materials and Methods

### *Drosophila* Stocks

Stocks were obtained from the Bloomington Drosophila Stock Center or the Vienna Drosophila RNAi Center, unless otherwise noted. Identifying stock numbers are referenced in the figure legends. UAS-NOS was provided by Pat O’Farrell (Yakubovich *et al*. 2010). y,w; phm-GAL4{51A2} was provided by Alexander Shingleton (Mirth *et al*. 2005). elav-Gal80 was provided by Yuh Nung and Lilly Jan. hs-NOS ^Mac^ and UAS-NOS^IR-X^ was provided by Henry Krause (Cáceres *et al*. 2011). PTTH-GAL4 was provided by Michael O’Connor (McBrayer *et al*. 2007; Halme *et al*. 2010). UAS-dilp8::3xFLAG was provided by Maria Dominguez (Garelli *et al*. 2012).

**Figure 1:**
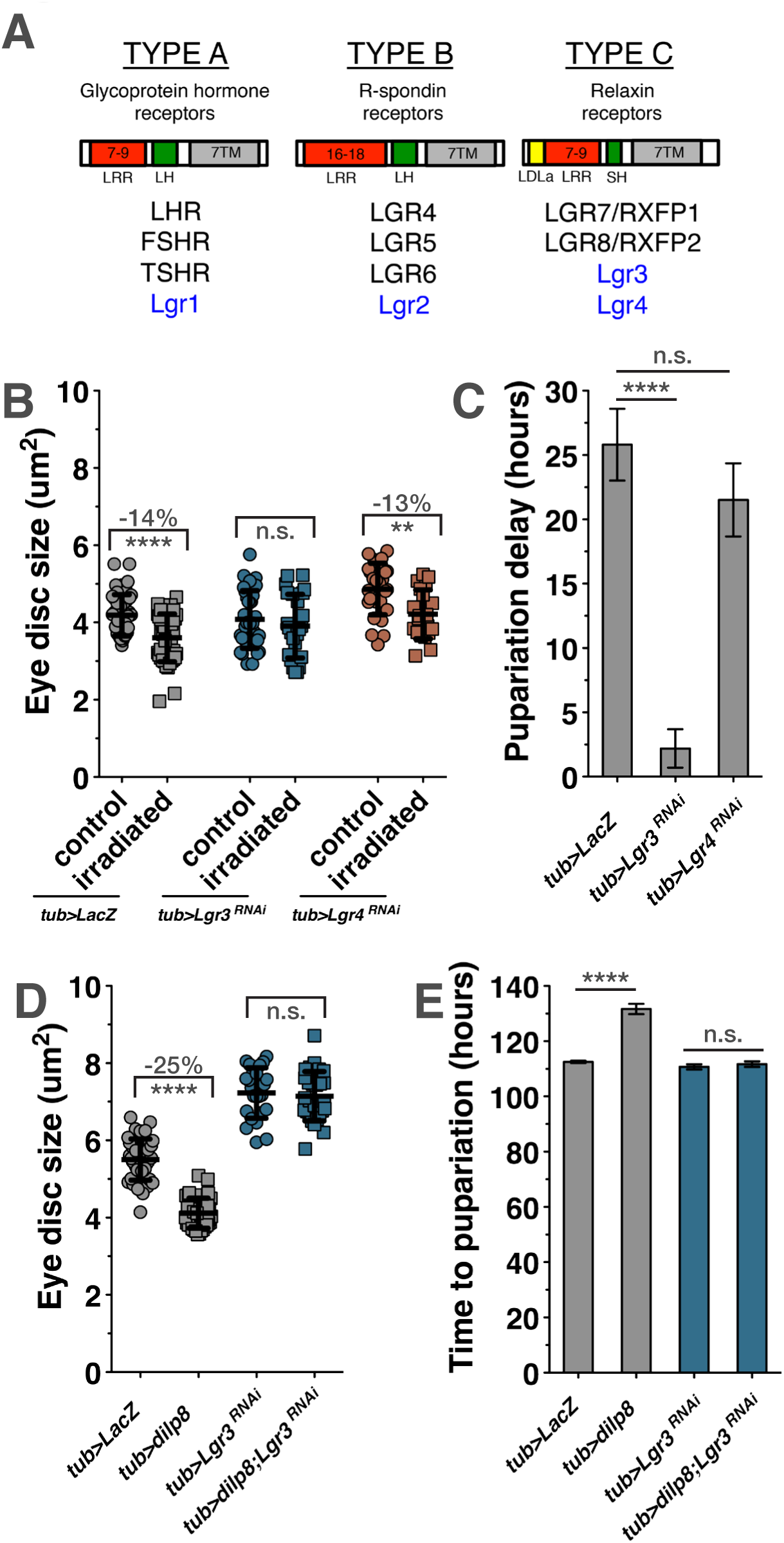
The *Drosophila* relaxin receptor homolog Lgr3 regulates Dilp8 mediated growth coordination and developmental delay during the regeneration checkpoint. (A) Comparison of the mammalian (black) and *Drosophila melanogaster* (blue) LGR protein types. LRR: Leucine-rich repeat domain – the number above denotes the number of repeats typically found among receptors of that LGR type; LH: long hinge domain; SH: short hinge domain; 7TM: seven transmembrane domain (B) Targeted irradiation to the posterior of the larva inhibits growth of the anterior-undamaged eye imaginal discs (*tub>LacZ*, irradiated vs control). Systemic expression of Lgr3-RNAi (*tub>Lgr3^RNAi^*) rescues growth restriction. Systemic expression of Lgr4-RNAi does not rescue growth restriction. (C) Full irradiation induces a developmental delay (*tub>LacZ*), which is rescued by systemic expression of Lgr3-RNAi. (D and E) Systemic expression of Dilp8 is sufficient to inhibit imaginal disc growth and developmental delay (*tub>dilp8*). Systemic expression of Lgr3-RNAi simultaneously with Dilp8 blocks both growth inhibition and Dilp8-induced delay (*tub>dilp8; Lgr3^RNAi^*). Growth: mean +/− SD. Time: mean of duplicate or triplicate experiments +/− SEM.** p<0.01, ****p<0.001 calculated by two-tailed Student’s t-test. See also Figure S1.

### Drosophila culture and media

Larvae were reared at 25° on standard Bloomington “Cornmeal, Molasses and Yeast Medium” supplemented with live bakers yeast granules. Developmental timing was synchronized through the collection of eggs during a 4-hour interval on grape agar plates. Twenty first-instar larvae were transferred to vials containing media 24hrs after egg deposition (AED).

### Targeted irradiation damage

Targeted irradiation experiments were conducted as previously described (Jaszczak *et al*. 2015). At 80hrs AED, shielded and unirradiated control larvae were immobilized on chilled glass cover slips and kept on ice during the duration of the irradiation. Ionizing irradiation was targeted to posterior portions of the larvae by placing a 0.5 cm^2^ strip of lead tape (Gamma) over the estimated anterior third of the larval body. Larvae were exposed to 25 Gy X-irradiation generated from a Faxitron RX-650 operating at 130kV and 5.0mA. Irradiated and control larvae were returned to cornmeal-molasses food and raised at 25° until dissection at 104hrs AED. Developmental delay after irradiation was assessed as previously described (Halme *et al*. 2010). Staged larvae were raised in petri dishes on standard media and irradiated in the food at 80 hrs AED.

### DAF2-DA assay

NO production was detected by 4,5-Diaminofluorescein diacetate (DAF2-DA, Sigma). Brain complexes were dissected in PBS, incubated in 10uM DAF2-DA for 10min at 28°, rinsed in PBS, fixed with 2-4% paraformaldehyde along with DAPI stain at 1:1000, rinsed in PBS, and imaged by confocal microscopy. DAF2-DA fluorescence was quantified in ImageJ (NIH) by measuring the mean gray value of each lobe of the PG normalized to the background fluorescence of the adjacent brain hemisphere. Fold change was calculated relative to the mean of the control for each genotype.

### Measurement of growth parameters

Time to pupariation, the time at which half the population had pupated, was calculated by recording the number of pupariated individuals every 12hrs. Developmental delay was calculated as time to pupariation of the experimental minus the control time to pupariation. Imaginal tissue area was measured using ImageJ on tissues dissected in phosphate-buffered saline (PBS), fixed in 4% paraformaldehyde, mounted in glycerol, and viewed by DIC on a Zeiss Axioplan2 microscope.

### X-gal staining

Tissues were dissected in PBS and fixed for 15min in 1% gluteraldehyde, incubated at 4^o^ overnight, rinsed in PBS, and mounted in glycerol.

### Results and Discussion

The *Drosophila* relaxin receptor homolog, Lgr3, is required for growth coordination and delay during the regeneration checkpoint

Based on the structural similarities between Dilp8 and relaxin proteins, we sought to determine whether Dilp8 activity is dependent on a *Drosophila* relaxin receptor homolog. *Drosophila* has four LGR proteins, of which only Lgr3 and Lgr4 share structural homology with the type C relaxin receptors (Fig. 1A) (Van Hiel *et al*. 2014). Lgr3 and Lgr4 have recently been shown to be expressed in many tissues throughout larval development (Van Hiel *et al*. 2014). To test whether these *Drosophila* relaxin homologs are necessary for growth coordination or developmental delay during the regeneration checkpoint, we ubiquitously expressed UAS-driven RNAi transgenes against each of the two receptors using *tubulin-Gal4*. We then activated the regeneration checkpoint in these larvae through targeted irradiation, producing damage in posterior tissues of the larvae while protecting anterior tissues like the eye imaginal discs and the PG (see Experimental Procedures and (Jaszczak *et al*. 2015)). Following posterior irradiation, the growth of anterior tissues is normally reduced due to Dilp8-dependent growth coordination (Jaszczak *et al*. 2015). RNAi inhibition of *Lgr3*, but not *Lgr4*, reduces checkpoint growth inhibition, restoring the growth of undamaged tissues in larvae with targeted irradiation (Fig. 1B), and also reduces checkpoint delay (Fig. 1C). This was confirmed with a second Lgr3-targeting RNAi transgene (JF03217), as well a third RNAi-expressing line that targets distinct sequences in *Lgr3* (HMC04196) (Fig. S1A,B). Additionally we tested

RNAi targeted to the other *Drosophila* LGR genes and found that neither *Lgr1* nor *Lgr2* depletion reduced damage-induced growth inhibition (Fig. S1C) or developmental delay (Fig. S1D), suggesting that they do not mediate Dilp8 activity. However, we did observe that expression of either *Lgr1* or *Lgr2* RNAi produced a significantly longer delay following irradiation than in control larvae (Fig. S1D). Therefore, these genes may play other roles in the regulation of developmental timing.

Expression of Dilp8 alone, in the absence of damage, is sufficient to induce growth restriction and developmental delay (Fig1D,E) (Colombani *et al*. 2012; Garelli *et al*. 2012; Jaszczak *et al*. 2015). To test whether Dilp8 depends on Lgr3 for these activities, we co-expressed Dilp8 and an RNAi targeting *Lgr3* using the *tubulin-Gal4* driver. In larvae depleted of *Lgr3*, Dilp8-induced growth inhibition and developmental delay were both rescued (Fig. 1D,E). Together, these data demonstrate that of the *Drosophila* LGR proteins, Lgr3 alone is required for Dilp8-dependent coordination of growth and developmental delay during the regeneration checkpoint.

### Lgr3 mediates Dilp8 activation of NOS in the PG and is necessary for growth coordination during the regeneration checkpoint

To identify tissues where Lgr3 is expressed and thus may respond to Dilp8 signaling, we used a collection of Lgr3 enhancer-Gal4 transgenes (Pfeiffer *et al*. 2008)(Fig. S2A). These transgenes allow us to express nuclear-localized β-galactosidase in tissues where Lgr3 regulatory regions are transcriptionally active. Following staining, we observe that these enhancer-Gal4 transgenes express predominantly in the central nervous system (CNS) (Fig. S2B-F). Additionally, the enhancer-Gal4 transgene 18A01 consistently expresses in both the CNS and PG (Fig 2A and S2E, G). All PGs analyzed expressed the 18A01 transgene, however the expression was often only observed in a subset of PG cells. This pattern of expression suggests the activity of the 18A01 regulatory region in the PG could be dynamic. An overlapping enhancer region, 17H01 also produced a minority of PG tissues where expression could be observed in a single cell (Fig. S2G). Since none of the other transgenes tested produced any detectable expression in the PG, we concluded that the PG expression observed in 18A01 and 17H01 was specific to these enhancer elements.

**Figure 2:**
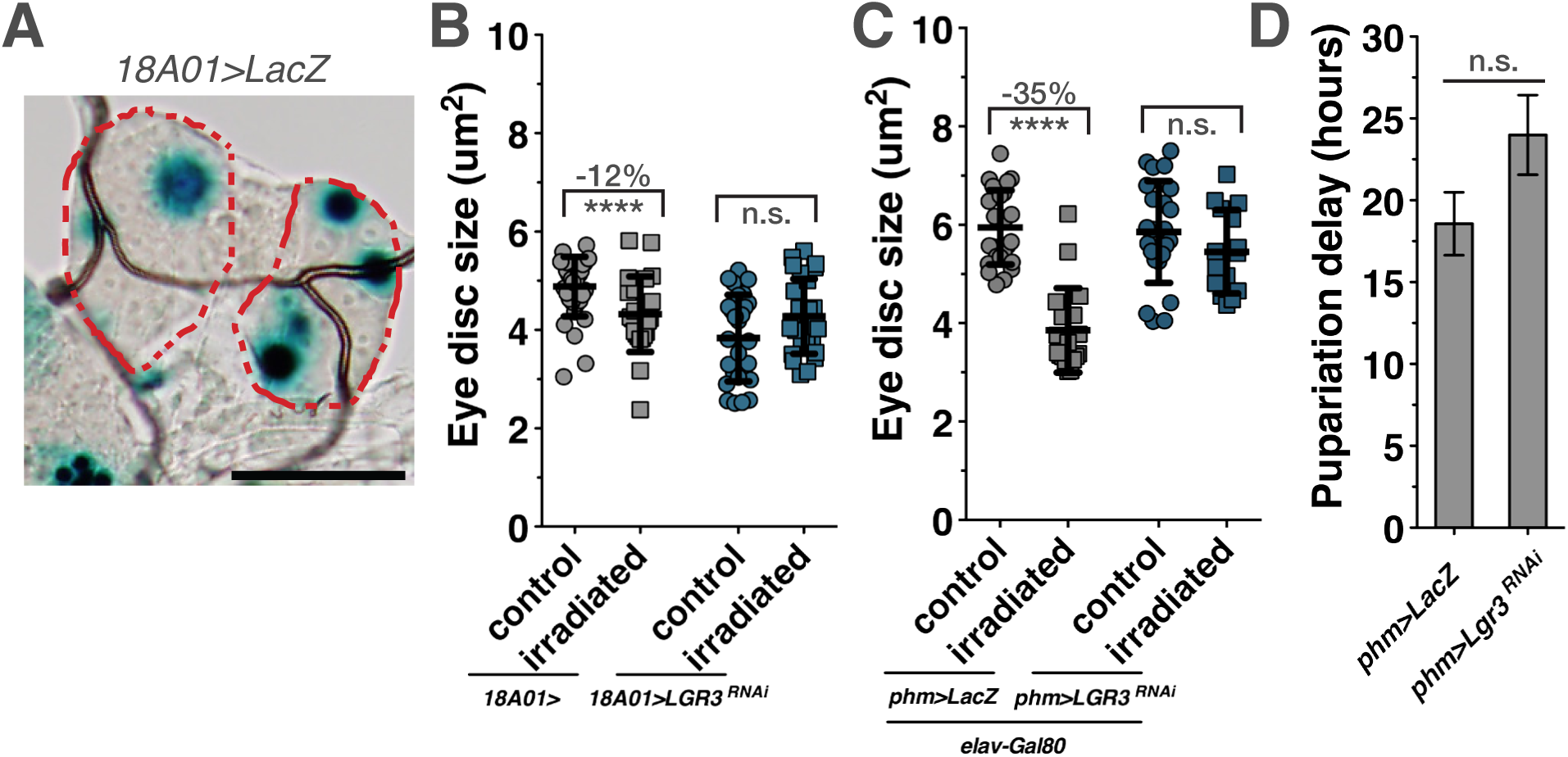
Lgr3 in the PG regulates growth coordination during the regeneration checkpoint. (A) Expression of nuclear-localized β-galactosidase in the PG visualized with X-gal staining in 104hr AED larva driven by enhancer 18A01 (*18A01>LacZ*). PG outlined by red dashes. Scale bar = 50um. (B) Expression of Lgr3-RNAi with the Lgr3 enhancer-Gal4 (*18A01>Lgr3^RNAi^*) reduces growth inhibition induced by targeted irradiation. (C) Expression of Lgr3-RNAi in the PG while also expressing the Gal4 inhibitor Gal80 in neurons (*elav-Gal80, phm>Lgr3^RNAi^*) rescues growth inhibition induced by targeted irradiation. (D) Expression of Lgr3-RNAi in the PG does significantly not affect developmental delay induced by irradiation. Growth: mean +/− SD. Time: mean of triplicate experiments +/− SEM. ****p<0.001 calculated by two-tailed Student’s t-test. See also Figure S2.

We have recently reported that Dilp8 coordinates growth through the activation of NOS in the PG (Jaszczak *et al*. 2015), therefore we tested whether Lgr3 is required for growth regulation in the cells that express the 18A01 enhancer-Gal4 transgene. When an *Lgr3-targeting* RNAi was expressed using the 18A01 enhancer-Gal4, growth inhibition of the undamaged imaginal discs does not occur (Fig. 2B), suggesting that the 18A01 enhancer expresses in cells that require Lgr3 to produce growth coordination following damage. To determine whether Lgr3 activity was specifically required in the PG for growth coordination following damage, we examined growth coordination in larvae expressing *Lgr3-*RNAi using the PG-specific *phantom-Gal4* (Mirth *et al*. 2005) driver. To ensure that we were exclusively assessing the role of Lgr3 in the PG, we also included a neuron-expressed Gal4 repressor (*elav-Gal80*). In these larvae, we observed that growth inhibition of undamaged imaginal discs was substantially reduced when compared to control larvae (Fig. 2C). These results demonstrate that Lgr3 activity in the PG is necessary for growth coordination following regeneration checkpoint activation. We also observed that RNAi depletion of *Lgr3* in the PG has no effect on the developmental delay produced by activation of the regeneration checkpoint (Fig. S3A). This observation is consistent with what we have reported for NOS activity, where NOS activation in the PG is necessary for damage and Dilp8-mediated growth inhibition, but not developmental delay (Jaszczak *et al*. 2015). Therefore, we speculated that Lgr3 might be regulating NOS activity in the PG during the regeneration checkpoint.

To determine whether PG expression of Lgr3 is required for the damage-induced NOS activity, we used the fluorescent reporter molecule 4,5-diaminofluorescein diacetate (DAF2-DA) to measure NOS activity through NO production in the PG. Using this assay, we have previously shown that Dilp8 expression is sufficient to induce NOS activation in the PG (Jaszczak *et al*. 2015). After posterior irradiation of larvae, NO production increases in the PG in a Dilp8 dependent manner (Fig. 3A and B). When we express an Lgr3-targeting RNAi in the PG with the *phantom-Gal4* driver, activation of NOS is no longer detected in the PG following irradiation (Fig. 3C). These data demonstrate that Lgr3 activity in the PG is required for NOS activation during the regeneration checkpoint. We have previously shown that NOS is required for Dilp8-mediated growth inhibition (Jaszczak *et al*. 2015). To establish that NOS functions downstream of Lgr3, we determined whether artificially increasing NOS activity could restrict growth independently of Lgr3 function in the PG. To do this, we overexpressed NOS along with the Lgr3-targeting RNAi in the PG using *phantom-Gal4*. We found that even when Lgr3 is depleted from the PG, NOS is still able to inhibit imaginal disc growth (Fig. 3D). Together, these data demonstrate that Lgr3 in the PG functions upstream of NOS, is necessary for NOS activation, and is required for Dilp8-mediated growth control through NOS.

**Figure 3:**
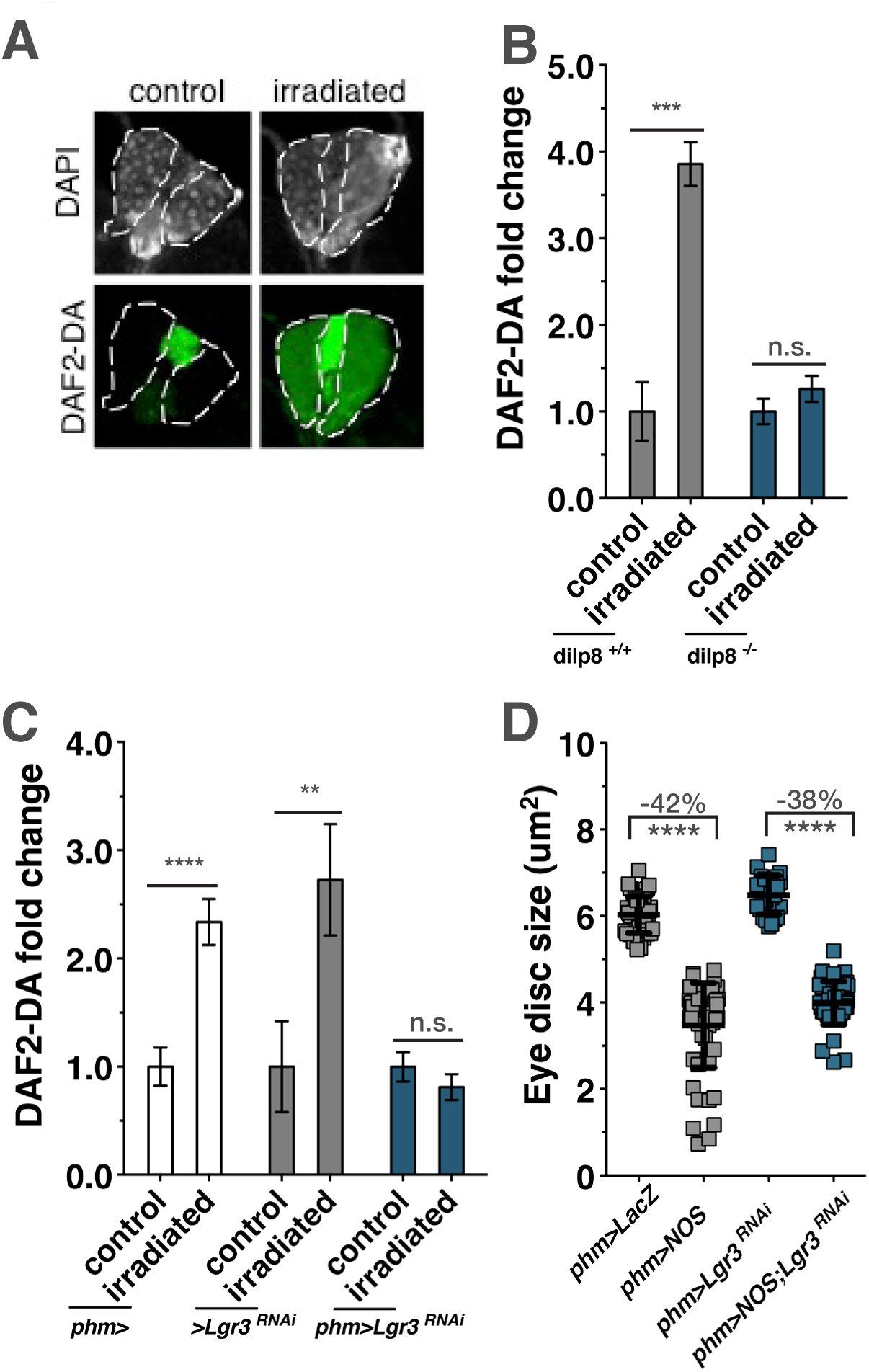
Lgr3 in the PG regulates NOS activity during the regeneration checkpoint. (A) Targeted irradiation increases NO production in the PG. (gray: DAPI, green: DAF2-DA). (B) Activation of NO production in the PG after targeted irradiation is lost in larva mutant for Dilp8 (n = 5–10 PGs) (C) Expression of Lgr3-RNAi in the PG blocks activation of NO production after targeted irradiation. (n = 5–10 PGs) (D) Overexpression of NOS in the PG (*phm>NOS*) inhibits imaginal disc growth even when Lgr3-RNAi is also expressed (*phm>NOS;Lgr3^RNAi^*). Growth: mean +/− SD. +/− SEM. * p<0.05, ***p<0.005, ****p<0.001 calculated by two-tailed Student’s t-test.

### Neuronal Lgr3 activity regulates regeneration checkpoint delay and growth coordination

Since all the Lgr3 enhancer-Gal4 transgenes analyzed express in the CNS (Fig. S2B-F), we wanted to determine if Lgr3 activity in neurons is important for regulating systemic responses to damage during the regeneration checkpoint. In particular, Lgr3 function is essential for developmental delay in response to imaginal disc damage (Fig. 1C), but not through its activity in the PG (Fig. 2D). To test the neuronal function of Lgr3, we examined larvae that expressed Lgr3 under the control of the neuron-specific *elav-Gal4* driver. In *elav>Lgr3^RNAi^* larvae, irradiation damage produced essentially no delay in development (Fig. 4A), demonstrating that damage-induced Dilp8 requires Lgr3 function in the brain to regulate developmental timing. Unexpectedly, depletion of *Lgr3* in neurons also completely eliminated growth coordination following targeted irradiation (Fig. 4B). This observation was confirmed with the neuron-specific *synaptobrevin-Gal4* (Pauli *et al*. 2008) expression of Lgr3-targeted RNAi, which also eliminated growth coordination following targeted irradiation (Fig. S3A). However, *Lgr3-*targeted RNAi in glial cells using *repo-Gal4* did not rescue growth inhibition or developmental delay (Fig. S3B,C) demonstrating that Lgr3 function is required specifically in neurons for growth coordination during the regeneration checkpoint.

**Figure 4:**
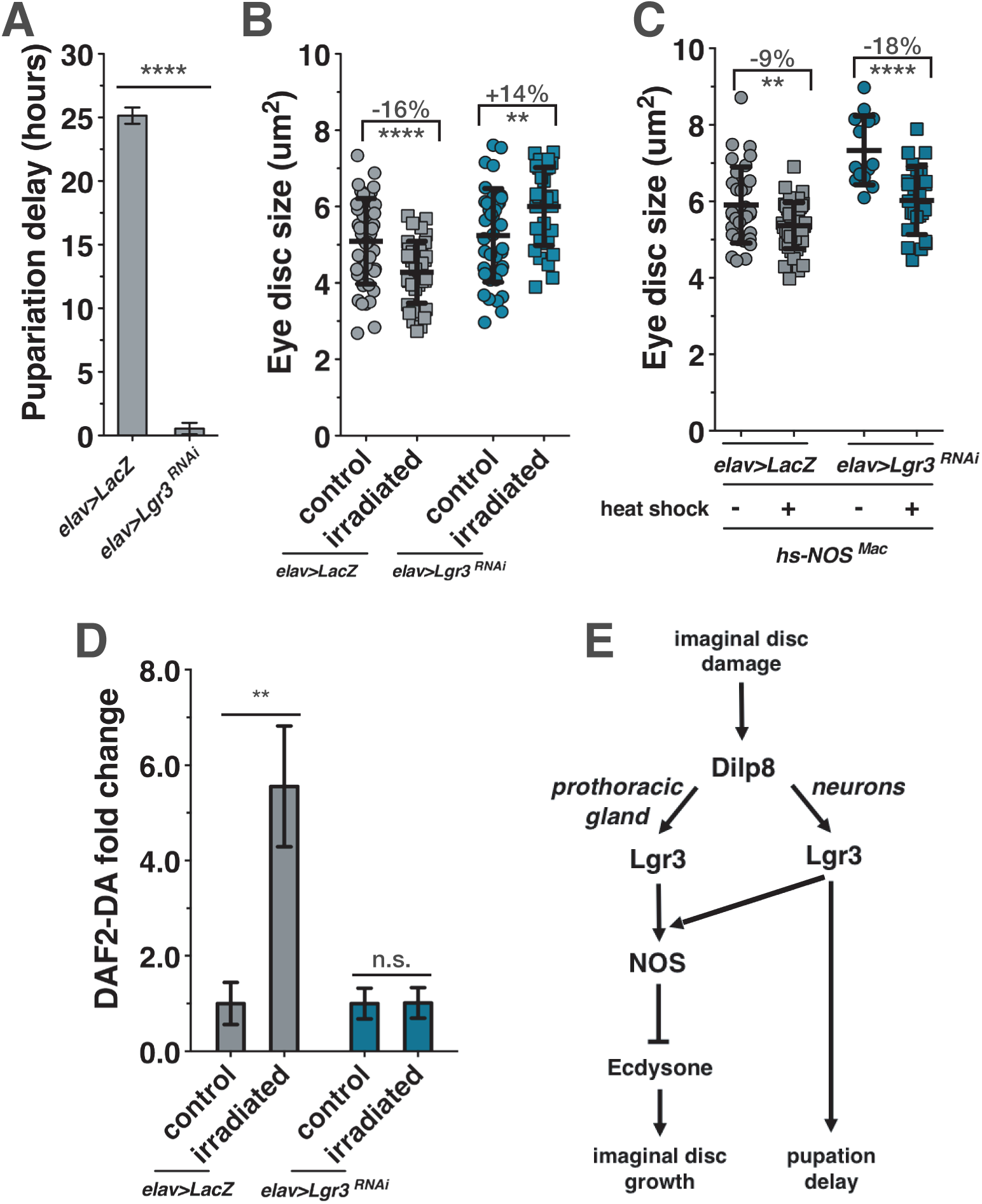
Lgr3 in neurons regulates developmental delay and also regulates growth coordination during the regeneration checkpoint through NOS activity. (A) Expression of Lgr3-RNAi in neurons (*elav>Lgr3^RNAi^*) largely abrogates developmental delay induced by irradiation. (B) Targeted irradiation of larvae expressing Lgr3-RNAi in neurons (*elav>Lgr3^RNAi^*) increases imaginal disc growth in contrast to the growth inhibition in the control (*elav>LacZ*). (C) Expression of Lgr3-RNAi in neurons (*elav>Lgr3^RNAi^*) does not block NOS inhibition of imaginal disc growth. NOS ^Mac^ was misexpressed by heat shock activation at 80hrs AED for 40 min in a 37° water bath. (D) Expression of Lgr3-RNAi in neurons blocks activation of NO production after targeted irradiation. (n = 5–10 PGs) (E) Lgr3 mediates growth coordination and developmental delay during the regeneration checkpoint through distinct tissues. Lgr3 in the PG regulates growth coordination, but not delay, through activation of NOS, which reduces ecdysone production. Lgr3 in the neurons mediates Dilp8 activation of developmental delay and also regulates growth coordination through regulation of NOS activity in the PG. Growth: mean +/− SD. Time: mean of duplicate or triplicate experiments +/− SEM. ** p<0.01, ****p<0.001 calculated by two-tailed Student’s t-test. See also Figure S3.

Regeneration checkpoint delay is the result of delayed expression of the neuropeptide PTTH (Halme *et al*. 2010), therefore we tested whether Lgr3 might be acting in the PTTH-expressing neurons (McBrayer *et al*. 2007) to directly regulate delay or growth inhibition. However, neither growth nor delay was affected by *Lgr3*-targeted RNAi expression specifically in the PTTH-expressing neurons (Fig. S3D,E). Therefore, other neurons expressing Lgr3 are likely communicating regeneration checkpoint activation to the PTTH-expressing neurons.

Since the *Lgr3*-dependent activation of NOS in the PG is required for growth coordination, we also tested whether NOS is required in the neurons for regulating *Lgr3*-dependent growth coordination and developmental delay during the regeneration checkpoint. Using a NOS-directed RNAi (Jaszczak *et al*. 2015) expressed in neurons (*elav>NOS^RNAi^*) during targeted irradiation, we found that neuronal depletion of *NOS* did not restore growth to undamaged tissues (Fig. S3F) or reduce developmental delay (Fig. S3G). This suggests that Lgr3 in neurons regulates growth through distinct cellular pathways from Lgr3 in the PG.

Together, these data indicate that Lgr3 is required: 1) in the CNS to mediate the effect of Dilp8 on developmental timing, and 2) in <u>both</u> the CNS and the PG to mediate Dilp8 effects on imaginal disc growth. To understand the relationship between these two roles for Lgr3 in regulating growth, we first sought to determine whether Lgr3 in the CNS is required for growth inhibition by NOS activation. To do this we used the heat shock promoter to overexpress NOS, which inhibits imaginal disc growth by reducing ecdysone production from the PG (Jaszczak *et al*. 2015), while also targeting expression of the *Lgr3* RNAi to neurons. We found that Lgr3 depletion from neurons has no effect on the ability of NOS to inhibit imaginal disc growth (Fig. 4C), demonstrating that NOS functions downstream of Lgr3 in the CNS. We then tested whether CNS Lgr3 functions upstream of NOS to regulate growth. We could determine this by examining the activation of NOS following damage in larvae where CNS expression of Lgr3 is depleted. To do this, we measured NO production in the PG with the fluorescent reporter DAF2-DA following irradiation damage in control and *elav>Lgr3^RNAi^* larvae. After targeted irradiation of larvae, we found that NO production did not increase in the PG when *Lgr3*-RNAi expression is targeted to the neurons (Fig. 4D). This demonstrates that neuronal Lgr3 functions upstream of NOS and regulates the ability of NOS to be activated in the PG. Therefore, Lgr3 in the CNS and in the PG are both required for the activation of NOS to mediate Dilp8 regulation of imaginal disc growth.

Our observations demonstrate that the *Drosophila* relaxin receptor Lgr3 mediates the effect of Dilp8 on developmental timing and growth coordination during *Drosophila* imaginal disc regeneration (Fig. 4E). In three recently published parallel studies, researchers have demonstrated that Lgr3 is required in a specific subsets of neurons in the CNS to coordinate the effects of Dilp8 on growth and developmental timing (Colombani *et al*. 2015; Garelli *et al*. 2015; Vallejo *et al*. 2015). This published work is consistent with our findings that neuronal disruption of Lgr3 expression is required for growth regulation and developmental delay. Our study here complements and extends these findings by demonstrating: 1) that the role of Lgr3 in growth regulation and developmental delay are separable through Lgr3 function in the PG, 2) that growth regulation depends on both Lgr3 activity in the CNS <u>and</u> the PG, and 3) that Lgr3 function in the CNS and in the PG is required for damage-induced NOS activation in the PG, explaining how Lgr3 function in both these two tissues is necessary for growth coordination.

Previous understanding of the biological activities of relaxins and their receptors have been largely restricted to their roles in sexual development and the function of the reproductive organs (Bathgate *et al*. 2013). We demonstrate that *Drosophila* relaxin receptor Lgr3 is necessary for coordinating growth between tissues during a regeneration checkpoint. Recently, allele polymorphisms at Lgr8/RXFP2 (the mammalian homologue of the *Drosophila* Lgr3) has been demonstrated to be an important genetic determinant of relative horn size within a population of wild Soay sheep (Johnston *et al*. 2013). This suggests a role for relaxin receptors in regulating growth and organ allometry is likely to be conserved in mammals.

## Acknowledgements

This work was funded in part by March of Dimes Basil O’Connor Starter Scholar award to AH (#5-FY12-60), and the National Institutes of Health to AH (RO1GM099803) and to JSJ (T32GM008136). We thank M. Dominguez, A. Shingleton, P. O’Farrell, H. Krause, and M. O’Connor for *Drosophila* stocks. We thank P. Leopold for helpful conversations on the work in this manuscript; and D. Castle, D. DeSimone, and D.G. Halme, for suggestions on the manuscript.

## Genotypes

Figure 1:

(B,C)

*UAS-dicer2/+; tub-GAL4/UAS-LacZ.NZ* (*tub-Gal4 line is derived from BL5138*)

*UAS-dicer2/+; tub-GAL4/UAS-GL01056*

*UAS-dicer2/+; tub-GAL4/UAS-JF03070*.

*UAS-LacZ.NZ* BL3956.

(D,E)

*UAS-dicer2/+; tub-GAL4/UAS-LacZ. NZ*

*UAS-dicer2/ UAS-dilp8::3xFLAG; tub-GAL4/+*

*UAS-dicer2/+; tub-GAL4/UAS-GL01056*

*UAS-dicer2/ UAS-dilp8::3xFLAG; tub-GAL4/ UAS-GL01056*

Figure 2:

(A)

*18A01-GAL4/UAS-LacZ. NZ*

(B)

*18A01-GAL4/+] [18A01-GAL4/UAS-GL01056*

(C)

*phm-GAL4/+;elav-GAL80/ UAS-LacZ. NZ*

*[phm-GAL4/HMC04196;elav-GAL80/ +*

(D)

*phm-GAL4/+;UAS-dicer2/ UAS-LacZ. NZ*

*phm-GAL4/+;UAS-dicer2/ UAS-GL01056*

Figure 3:

(A)

*Bx*(*MS1096*)

(B)

*Bx*(*MS1096*)

*Bx*(*MS1096*)*;;dilp8MI00727*

(C)

*phm-GAL4;UAS-dicer2*

*UAS-GL01056/TM6B*

*phm-GAL4/+;UAS-dicer2/ UAS-GL01056*

(D)

*phm-GAL4/+;UAS-dicer2/ UAS-LacZ. NZ*

*phm-GAL4/UAS-NOS;UAS-dicer2/ UAS-GFP*

*phm-GAL4/+;UAS-dicer2/ UAS-GL01056*

*phm-GAL4/UAS-NOS;UAS-dicer2/ UAS-GL01056*

Figure 4:

(A,B,D)

*elav-GAL4/ UAS-LacZ.NZ*

*elav-GAL4/ UAS-GL01056*

(C)

*hsNOSmac, elav-GAL4/ UAS-LacZ.NZ*

*hsNOSmac,elav-GAL4/ UAS-GL01056*

Figure S1:

(A)

*UAS-dicer2/+;tub-GAL4/UAS-LacZ.NZ*

*UAS-dicer2/+;tub-GAL4/JF03217*

(B)

*UAS-dicer2/+;tub-GAL4/UAS-LacZ.NZ*

*UAS-dicer2/+;tub-GAL4/JF03217*

*UAS-dicer2/ HM04196;tub-GAL4/+*

(C,D)

*UAS-dicer2/+;tub-GAL4/UAS-LacZ.NZ*

*UAS-dicer2/+;tub-GAL4/ JF02659*

*UAS-dicer2/+;tub-GAL4/ JF02678*

Figure S2:

(B,G)

*17G11-GAL4/UAS-LacZ.NZ*

(C,G)

*18C07-GAL4/UAS-LacZ.NZ*

(D,G)

*17H01-GAL4/UAS-LacZ.NZ*

(E,G)

*18A01-GAL4/UAS-LacZ.NZ*

(F,G)

*19B09-GAL4/UAS-LacZ.NZ*

Figure S3:

(A,B,E)

*phm-GAL4/+;UAS-dicer2/ UAS-LacZ.NZ*

*phm-GAL4/+;UAS-dicer2/ UAS-GL01056*

(C)

*18A01-GAL4/+*

*18A01-GAL4/ UAS-GL01056*

(D)

*UAS-dicer2/+;tub-GAL4/UAS-LacZ.NZ*

*UAS-dicer2/+;tub-GAL4/UAS-GL01056*

(F)

*elav-GAL4/ UAS-LacZ.NZ*

*elav-GAL4/ UAS-GL01056*

Figure S4:

(A)

*syb-GAL4/+*

*syb-GAL4/ UAS-GL01056* (*syb-GAL4 BL51635*)

(B,C)

*UAS-dicer2/+;repo-GAL4/UAS-LacZ.NZ*

*UAS-dicer2/+;repo-GAL4/UAS-GL01056* (*repo-GAL4* BL7415)

(D,E)

*UAS-dicer2/+;PTTH-GAL4/UAS-LacZ.NZ*

*UAS-dicer2/+;PTTH-GAL4/UAS-GL01056*

(F)

*elav-GAL4/ UAS-LacZ.NZ*

*elav-GAL4/ NOS^IR-X^*

(G)

*NOS^IR-X^*

*elav-GAL4/ NOS^IR-X^*

## Supplemental Figures

**Figure S1:**
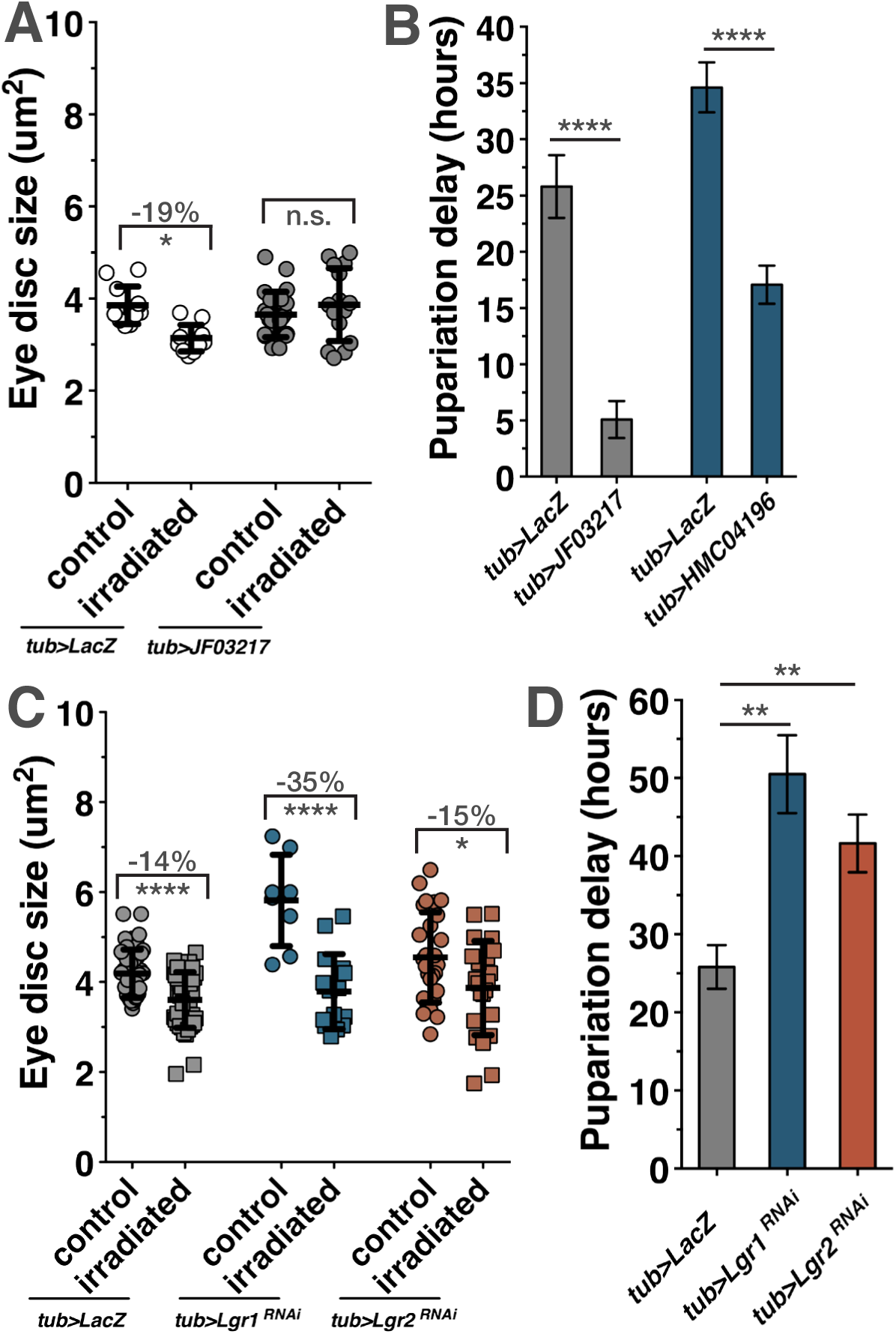
Related to Figure 1. LGR1 and LGR2 do not regulate growth coordination. (A and B) Systemic expression of Lgr3-RNAi rescues growth restriction induced by targeted irradiation and developmental delay induced by irradiation. (C and D) Systemic expression of Lgr1-RNAi or Lgr2-RNAi does not rescue growth restriction induced by targeted irradiation or developmental delay induced by irradiation. Growth: mean +/− SD. Time: mean of duplicate or triplicate experiments +/− SEM. * p<0.05, ****p<0.001 calculated by two-tailed Student’s t-test.

**Figure S2:**
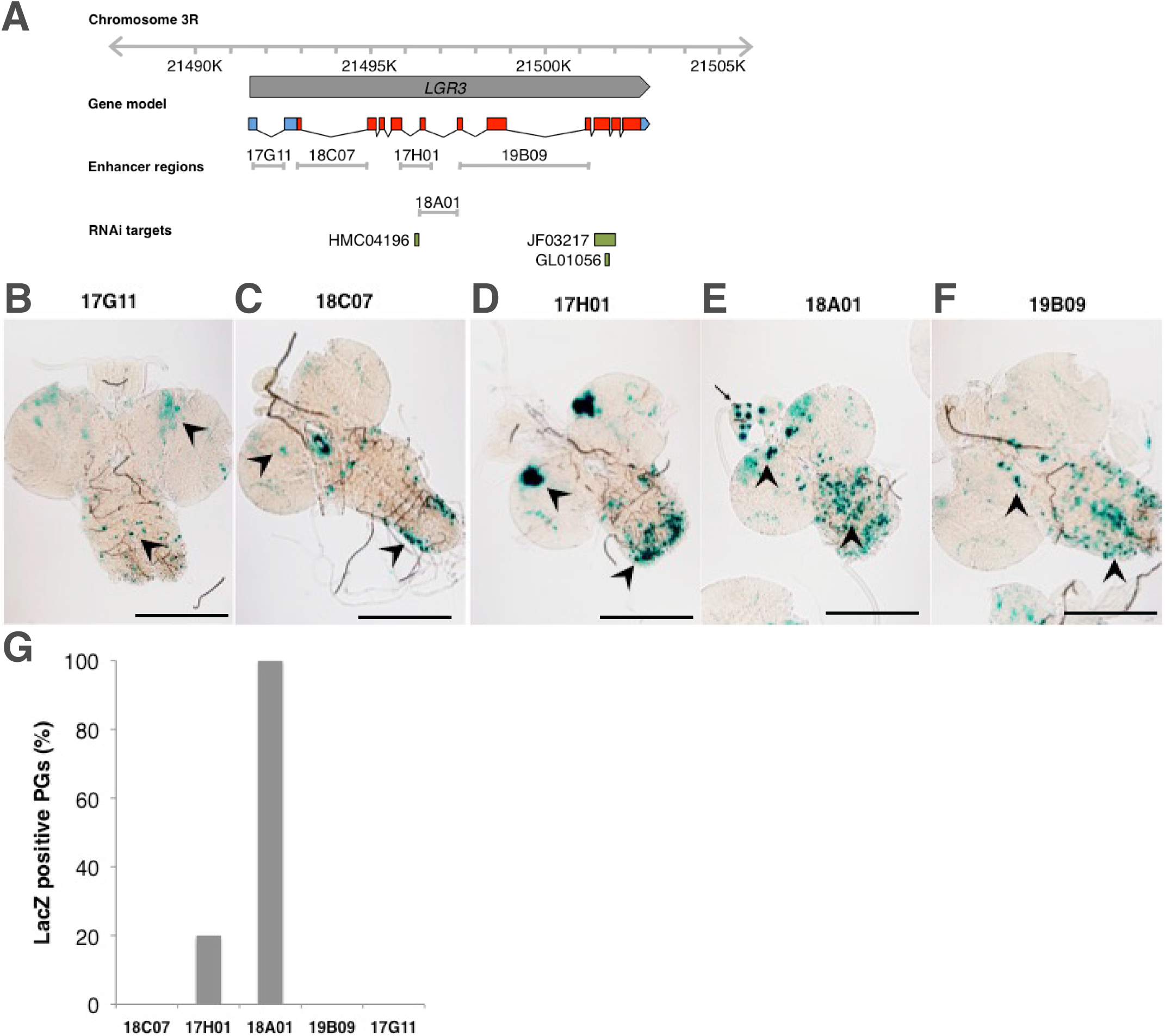
Related to Figure 2. Enhancer elements of Lgr3 express in the larval CNS and PG. (A) Gene map of Lgr3. Corresponding regions of enhancer elements used to generate enhancer-Gal4 transgenes. Lgr3 RNAi targeting regions. Blue boxes: 3’ and 5’ UTR. Red boxes: exons. Green boxes: RNAi target regions. (B-F) Expression of nuclear-localized β-galactosidase visualized by X-gal staining in 104hr AED larva. The arrow denotes enhancer activity observed in the PG (E). Arrowheads denote regions with recurring patterns of CNS enhancer activity. (G) Percent PGs with LacZ activity from Gal4-enhancer lines. 17H01 expression observed in 20% of isolated PGs only labeled a single cell. In contrast 18A01 enhancer consistently expressed in 4–10 cells of every PG. (n=8–11 PGs for each enhancer) Scale bars = 200um.

**Figure S3:**
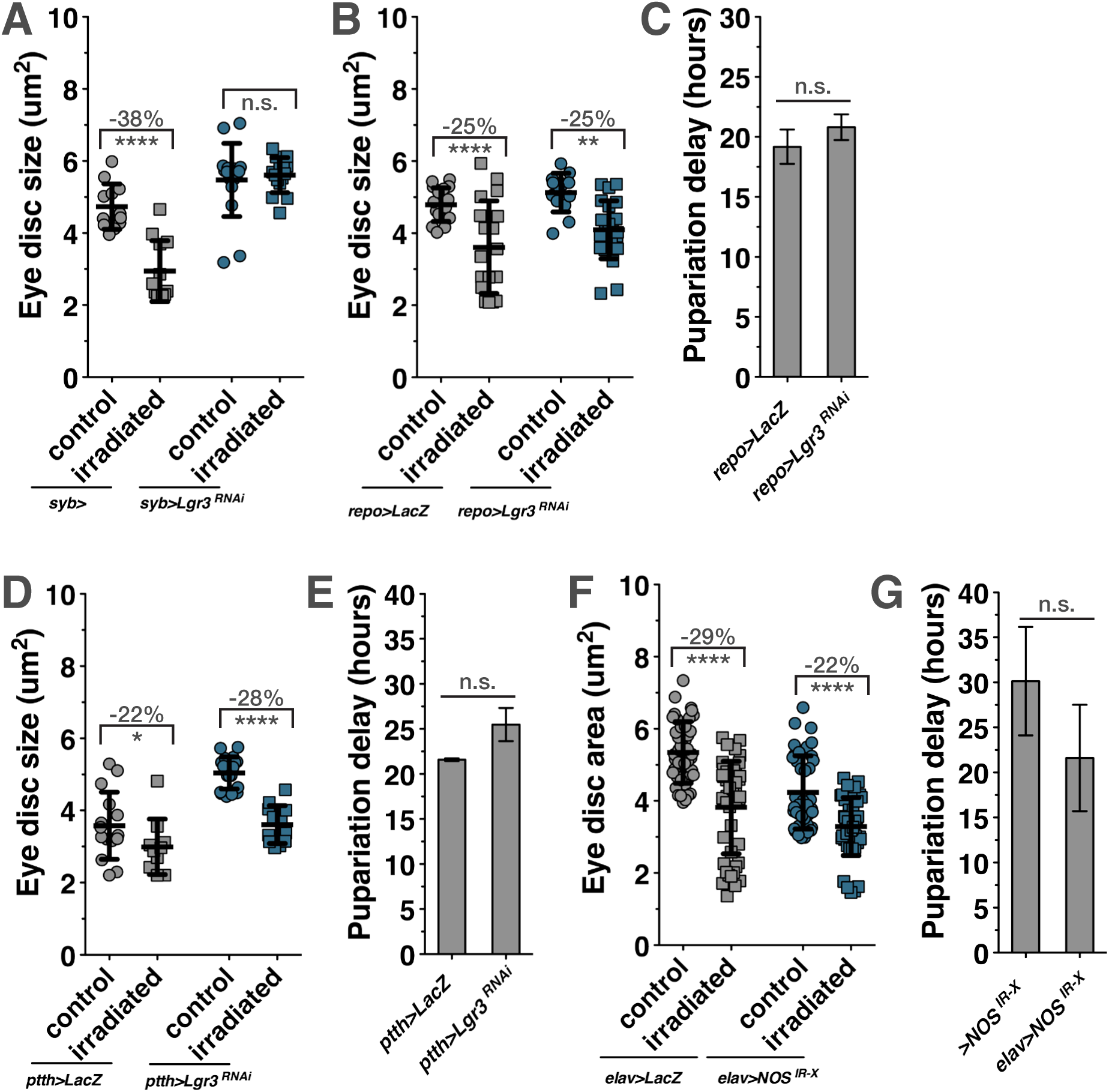
Related to Figure 4. Lgr3 in CNS neurons regulates growth and timing, but not by directly acting in PTTH neurons, or by CNS NOS activity. (A) Expression of Lgr3-RNAi with a neuronal-specific driver (*syb>Lgr3^RNAi^*) rescues growth inhibition induced by targeted irradiation. (B and C) Expression of Lgr3-RNAi with a glial-specific driver (*repo>Lgr3^RNAi^*) does not rescue growth inhibition induced by targeted irradiation or developmental delay. (D and E) Expression of Lgr3-RNAi in the PTTH neurons (*ptth>Lgr3^RNAi^*) does not rescue growth inhibition induced by targeted irradiation or developmental delay. (F and G) Expression of NOS-RNAi in neurons (*elav>NOS^IR-X^*) does not rescue growth inhibition induced by targeted irradiation or developmental delay. Growth: mean +/− SD. Time: mean of duplicate or triplicate experiments +/− SEM. * p<0.05, ** p<0.01, ****p<0.001 calculated by two-tailed Student’s t-test.

